# The cellular mammalian clock regulates *Staphylococcus aureus* invasion in epithelial cells

**DOI:** 10.1101/2025.05.05.652254

**Authors:** Pooja Agarwal, Giridhar Chandrasekharan, Sreshtha Nayek, Jaspreet Grewal, Robert Dallmann, Meera Unnikrishnan

## Abstract

An endogenous biological clock, the circadian clock, coordinates life with the 24-hour day/night environmental cycle. In mammals, a central pacemaker in the suprachiasmatic nucleus of the hypothalamus coordinates timing between peripheral clocks and with the environment and, for example, modulates immune responses to infections. However, its role in controlling bacterial infections at a cellular level is not understood. Here, we investigate the role of the host cellular clock during infection by a highly drug-resistant human pathogen, *Staphylococcus aureus.* Our findings revealed that *S. aureus* invasion into epithelial cells was dependent on the host circadian phase. Interestingly, cells deficient in BMAL1, a transcriptional activator and an essential clock protein, demonstrated increased bacterial uptake compared to parental A549 cells. The *BMAL1*-knockdown (KD) cells showed significant induction of GP340, a receptor of the *S. aureus* adhesin, SraP. An *S. aureus sraP* mutant did not exhibit rhythmic uptake into A549 cells or an increased uptake into *BMAL1*-KD compared to parental A549 cells. Of note, other bacterial adhesin mutants showed rhythmic and higher uptake in *BMAL1-*KD cells. Hence, we report that *S. aureus* epithelial cell invasion is clock-modulated and mediated through the *S. aureus* SraP-GP340 pathway, suggesting potential for host clock-directed therapy against this pathogen.

## Introduction

Circadian rhythms are intrinsic, about 24-hour cycles that govern a wide array of physiological processes in most living organisms, ranging from bacteria to humans^1,2^. These rhythms are regulated by a cell autonomous internal time-keeping system known as the circadian clock, which allows organisms to anticipate and adapt to daily environmental changes, synchronize metabolic functions, and maintain overall homeostasis^3^.

In mammals, a central clock in the hypothalamic suprachiasmatic nucleus (SCN) coordinates rhythms in all other organ, tissue and cellular clocks relative to the environmental light/dark cycles. The molecular basis of circadian rhythms involves coupled transcriptional-translational feedback loops consisting of more than a dozen core clock genes^1^, including the central positive and negative arm transcription factors BMAL1, CLOCK and PERIODs, CRYPOTCHROMEs, respectively. These genes interact to produce proteins that oscillate in abundance over an approximate 24-hour cycle, ultimately regulating downstream processes involved in metabolism, immunity, inflammation and various physiological responses^3^.

Recent research has underscored the significant influence of the circadian clock on immune regulation and host-pathogen interactions^4,5^. The synchronization between the circadian system and immune responses ensures that the body can efficiently combat pathogens at optimal times of the day^6^. For example, studies have shown that immune cells exhibit diurnal variations in activity, with certain populations peaking at specific times, thereby enhancing the body’s ability to respond to infections^7^. Disruptions to circadian regulation, whether due to genetic, environmental, or behavioural factors, can impair these defences, leading to increased susceptibility to infections and altered pathogen behaviour^8,9^. Additionally, dysregulation of the host circadian clock has been linked with increased susceptibility to bacterial infection^10^.

*Staphylococcus aureus*, a Gram-positive bacterium, is a highly drug-resistant opportunistic human pathogen responsible for a wide range of infections, from minor skin conditions to life-threatening diseases such as pneumonia, endocarditis, and sepsis^11,12^. *S aureus* is a facultative intracellular pathogen, which invades a range of mammalian cells, which offer a niche for replication, persistence and resistance to antimicrobials^13–16^. Recent findings suggest that disruption of circadian regulation can impair immune responses towards *S. aureus* infection^17^. However, at a cellular level, the molecular mechanisms underlying the role of the host clock in *S. aureus* infection remain unclear. Identifying host pathways involved may lead to the development of chronotherapeutic approaches that optimize timing for interventions against *S. aureus* infections.

In this study, we explore the role of the host circadian clock in modulating *S. aureus* infection of epithelial cells in an *in vitro* infection model. We demonstrate that *S. aureus* invasion is dependent on the mammalian circadian phases and that specific bacterial adhesins may control rhythmic invasion through interactions with clock-regulated host proteins. This study implicates potential molecular targets that could inform new strategies against *S. aureus* infection.

## Methods and Materials

### Bacterial strains, plasmids, and bacterial culture conditions

*Staphylococcus aureus* strains, *S. aureus* JE2 wild-type (WT) (BEI Resources), including *S. aureus* reporter strains (containing pMV158-mCherry or pMV158-GFP), *S. aureus*::mCherry (pOS1), *S. aureus*::GFP (pMV158), were cultured in tryptic soy broth (TSB, Merck) medium at 37°C with shaking, in the presence or absence of 5 µg/mL tetracycline, as appropriate.

Transposon mutants from the Nebraska transposon mutant library (https://ntml.unmc.edu) were obtained from BEI Resources for key adhesion factors, including FnbA (fibronectin-binding protein A) NR-46609, FnbB (fibronectin-binding protein B) NR-47271, and SraP (serine-rich adhesin for platelets) NR-46576. These mutants were cultured in Tryptic Soy broth (TSB) containing 5 µg/mL erythromycin as per instructions.

For infection experiments, bacterial cultures were initiated from frozen stocks one day before the assay and overnight cultures were prepared. On the day of the experiment, log-phase cultures were initiated by diluting the overnight cultures into fresh TSB medium. Bacteria were grown in a shaker at 37°C to an optical density at 600 nm (OD_600_) of 1.0, which is exponential growth phase for *S. aureus* JE2.

### Mammalian cell culture and circadian synchronization

Human lung epithelial adenocarcinoma A549 cells wild type (ATCC CCL-185) with or without sh*BMAL1* knockdown were cultured in high glucose DMEM supplemented with GlutaMAX (11584516, Gibco), 1% Penicillin/Streptomycin (P4458, Sigma-Aldrich) and dialysed 10% fetal bovine serum (FBS, 26400044, Gibco). Cells were maintained in a humidified atmosphere at 37°C with 5% CO₂. To evaluate *Staphylococcus aureus* invasion, mammalian cells were seeded in 24-well plates (662892, Greiner) or 4-chamber (177399, Nunc Lab-Tek) slides at a density of 3 × 10⁵ cells per well and incubated for 24 hours prior to infection.

For circadian synchronization experiments, cells were seeded at a density of 2 × 10⁵ cells per well and allowed to adhere for 24 hours. Synchronization was achieved using forskolin (FSK, 10 µM, S2449, Selleck Chemicals), a potent cyclic AMP (cAMP) activator known to reset the circadian clock of all cells in the culture vessel to a common phase^18,19^. Using this widely employed method, cellular clocks were reset 6, 12, 18, or 24 h before infection with *S. aureus* to assess circadian phase difference of bacterial invasion (Figure 1A). The time from FSK treatment is used as a phase-marker throughout.

**Figure 1.**
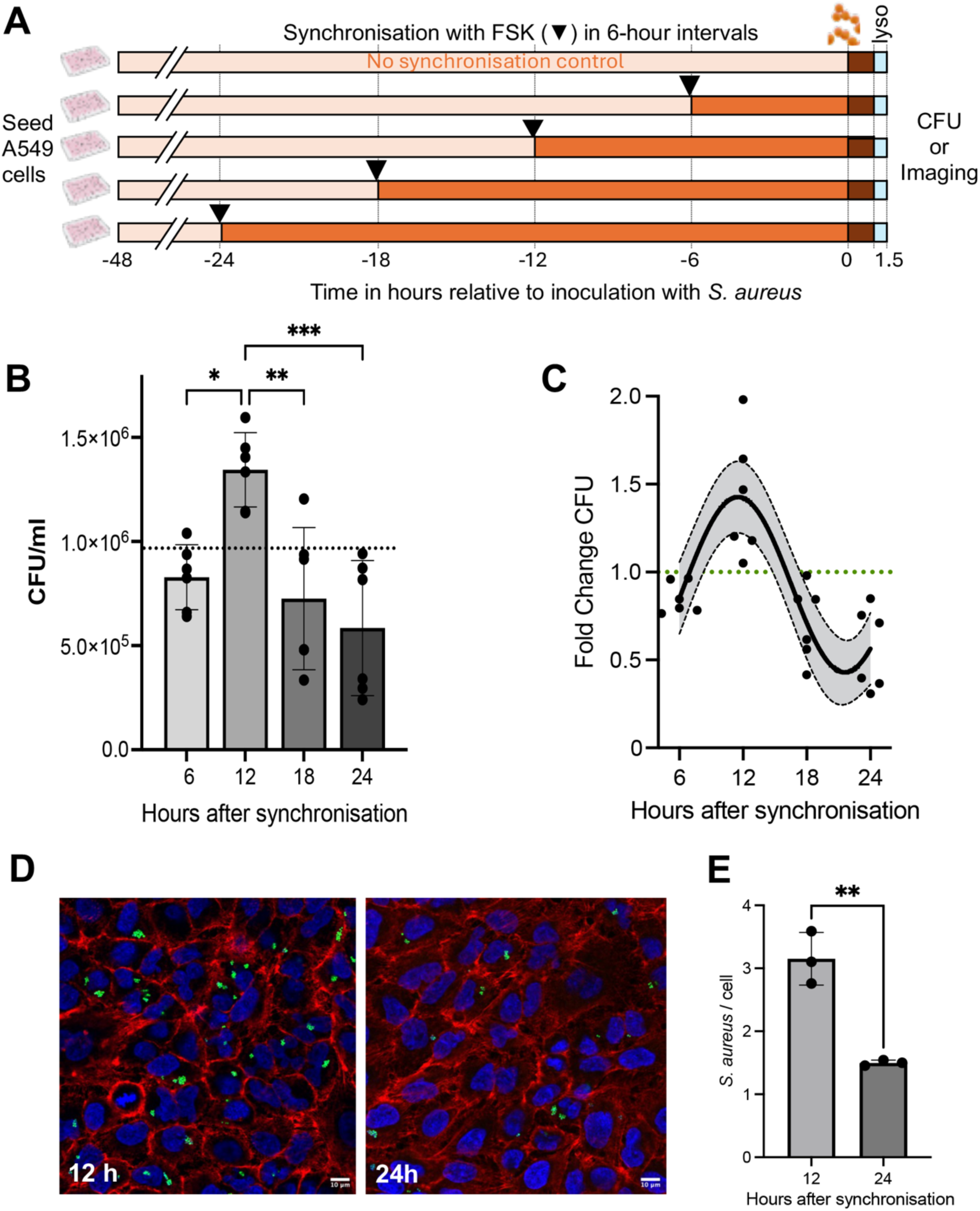
*S. aureus* invasion is modulated by epithelial cell circadian phase. **(A**) Schematic diagram showing experimental design. A549 cells were synchronized with forskolin (FSK) 24, 18, 12 and 6 h before infection with bacteria at an MOI of 10:1. FSK synchronized or unsynchronized cells were infected with wildtype USA300JE2 *S. aureus* for 1 h, and extracellular bacteria were removed by gentamicin/lysostaphin treatment. (**B**) Intracellular bacterial numbers determined by colony counts (CFU) from different times after synchronization (green dashed line shows value for mean unsynchronized cells), mean ± standard deviation (SD), * *P* = 0.014, ** *P* = 0.0031 and *** *P* = 0.004 by one way ANOVA with a Tukey’s multiple comparisons test, N = 6. (**C**) Data from B plotted as fold-change in CFU compared to an unsynchronized control showing COSINOR best fit with 95% confidence interval. (**D**) Confocal images of epithelial cells, synchronized by FSK for 24 h and 12 h, green - foci of intracellular *S. aureus*-GFP, Red - actin stained by Phalloidin Alexa fluor 647, Blue - nuclei stained by DAPI. Scale bar = 10 μm. A single Z-section is shown (**E**) Bacteria (green) and cell nuclei (blue) were counted manually and the relative number of bacteria to A549 cells was calculated from 3 independent experiments, mean ± SD, *P* = 0.003, unpaired t-test.

For microscopy-based assessments, cells were seeded either in imaging 24-well plates or on 4-chamber slides to facilitate imaging of *S. aureus* interactions with epithelial cells.

### Generation of transgenic A549 cell lines

*BMAL1* knockdown cells were generated by lentiviral transgenesis following a published strategy^20^. In short, lentiviral particles were generated by transfection of HEK293FT cells (R70007, Invitrogen) with a third generation lenti-viral backbone pGIPZ for *shBMAL1* packaging and capsid plasmids (pCMVR8.74 and pMD2.G, Addgene) using TransIT-LT reagent (Mirrus Bio) as described^21^. Supernatant containing viral particles was sterile filtered and frozen at −80°C until use. For lentiviral transduction, viral supernatant with 8 μg/mL polybrene (TR-1003G, Sigma-Aldrich) was added onto 60% confluent A549 cells, and incubated at 37°C for 6 h, after which cells were washed and incubated for 3 days in standard media. Afterwards, media was supplemented with 10 µg/ml puromycin to select infected cells. Co-expressed eGFP was assessed as further confirmation that cells had integrated the *shBMAL1* construct.

### Invasion and survival assays

To evaluate the invasion of A549 cells, *S. aureus* cultured to OD_600_ of 1.0 (logarithmic growth phase) were harvested by centrifugation at 5000 xg for 10 minutes and washed twice with phosphate-buffered saline (PBS). The bacterial pellet was resuspended in PBS and the required volume of this culture was added to the cell culture medium to prepare the inoculum at 3–6 × 10^6^ CFU/ml. This inoculum was added to epithelial cells (1 ml/well), for 1 h to achieve a multiplicity of infection (MOI) of 10:1. Inoculum bacterial counts and MOI’s were confirmed by bacterial colony counts and epithelial cell counts for each experiment. Following the infection period, extracellular bacteria were removed by incubating cells in culture medium containing gentamicin (50 µg/mL) and lysostaphin (20 µg/mL) for 30 minutes. The cells were then washed three times with PBS and lysed with cold water to release intracellular bacteria. To enumerate colony forming units (CFU), cell lysates were serially diluted and plated onto tryptic soy agar (TSA; Sigma). To monitor the intracellular growth kinetics of bacteria using microscopy, infected cells were incubated with 2 μg/mL lysostaphin. Imaging was performed using either a BioTek Cytation 5 Multimode Reader (Agilent) or a confocal microscope (Olympus FV3000).

Analysis of rhythmic patterns for synchronised cells infected with *S. aureus* was conducted on bacterial CFU or the fold-changes of internalised bacterial CFU/ml (bacterial invasion) relative to unsynchronised cells. A COSINOR model was used as previously described ^22^, using the equation Y=(Mesor+Amplitude*cos(((2*pi)*(X-Acrophase))/Period)) with constraining Period to over 20 h. The linear model was Y= YIntercept + X*Slope with no further constraint. Data was fit to COSINOR and linear models and an extra sum-of-squares F-test was run at *P* < 0.01 with the linear model as null hypothesis.

### RNA-Seq and data analysis

To harvest RNA, parental and *shBMAL1* A549 cells were seeded and synchronised by FSK as described above. RNA was extracted at the indicated times post synchronisation using the RNeasy Plus kit (74134, Qiagen). Briefly, the cells were washed thrice in PBS and lysed in the RLT buffer from the kit. Genomic DNA was removed from the lysate using a gDNA eliminator spin column from the kit. 1 volume of 70% ethanol was added to the flowthrough and loaded on a RNeasy spin column. The column was centrifuged at 8000 xg for 30 s. This was followed by subsequent washes with RW1 and RPE buffer. Excess ethanol from the column was removed by spinning the column at 13000 xg for 1 min. Twenty ml RNase free buffer was added to the column and spun at 8000 xg for 1 min. RNA was quantified using the Qubit assay. The RNA samples (30 ng/µl) were sent to Novogene for library preparation after poly(A) enrichment and subsequently sequenced using a pair-ended 150 bp kit on an Illumina NovaSeq. Sequencing depth was at least 15 million reads per sample. Gene count tables were generated using Kallisto on the BioJupies platform^23^. All further analysis was conducted using iDEP 2.0^24^. The DESeq2 algorithm was used for differential gene expression analysis with FDR cutoff of 0.01. For gene normalisation variance stabilising transformation (VST) was used for data in PCA, heatmap. Single gene expression levels are reported as log2CPM+1 values transformed with EdgeR^25,26^. Parametric Gene Set Enrichment Analysis (PGSEA) of KEGG pathways was used for pathway analysis with FDR cutoff of 0.0001^27^. RNA-Seq data are available at the ELIXIR database ArrayExpress under accession number E-MTAB-15036.

### Western blotting

To prepare cell lysates, synchronized and non-synchronised cells were lysed using RIPA buffer. Samples were separated on a 4-12% SDS-PAGE gels (Mini-PROTEAN TGX) and then transferred onto a nitrocellulose membrane (Bio-Rad) using wet transfer. The membrane was blocked for 1 hour with 5% non-fat milk and then incubated overnight at 4°C with 1 mg/ml anti-BMAL1 mAb (Abcam, 93806), diluted 1:2000. Membranes were washed three times with Tris buffered saline with 0,1% Tween-20 (TBS-T) and incubated with a 1:2000-diluted HRP-conjugated secondary goat anti-mouse antibody (Abcam, 205718) for 1 hour at room temperature (RT). After washing three times with TBS-T, the signal was detected using a Pierce ECL Western Blot Substrate (Thermo Scientific, 10005943) according to the manufacturer’s instructions.

The blot was then stripped using stripping buffer, washed with TBS-T, blocked again for 1 hour with TBS containing 5% non-fat milk, and incubated overnight at 4°C with 1 mg/ml anti-Actin mAb (Sigma A5316), diluted 1:2000. The next day, membranes were washed three times with TBS-T and incubated with a 1:2000-diluted HRP-conjugated secondary goat anti-mouse antibody (Abcam, ab8227) for 1 h at RT followed by detection with ECL substrate as above.

### Flow cytometry

We used flow cytometry to quantify GP340 protein expression in parental A549 cells from four 6-hourly samples taken at 6, 12, 18 and 24 h after FSK synchronisation as described above, and from unsynchronised cells to compare A549 parental with *shBMAL1* cells. For this, live cells were trypsinized, >95% viability confirmed, washed in PBS and density adjusted to 1×10^6^ cells/ml. After blocking of cells with 1% BSA in PBS for 30 min, they were pelleted and stained with anti-GP340 rabbit antibody (1:500, Thermofisher, PA5-115313) in 1% BSA in PBS and incubated at 4°C for 1 h in dark. Before and after staining with secondary anti-rabbit IgG Alexa Fluor 647 conjugated antibody (1:1000, Cell Signalling, 4414) for 1 h in the dark, cells were washed three times in PBS, resuspended, and 30,000 events were recorded in triplicate in a flow cytometer (BD Biosciences LSRFortessa) for analysis. The round-the-clock experiments were performed on two independent occasions in triplicate and results were normalised to the average GP340+ per sample and then pooled for COSINOR analysis as described above.

### Microscopy and image analysis

For bacterial visualization via confocal microscopy, cells were fixed with 4% paraformaldehyde (PFA) for 1 hour, followed by three washes with PBS. Actin was stained with Phalloidin 650 in the dark for 1 h. After staining, cells were washed three times with PBS, air-dried, and mounted using a mounting medium containing DAPI for nuclear staining. Stained samples were observed using an Olympus FV3000 Confocal Laser Scanning Microscope. Images were analysed with ImageJ using Fiji^28^. The imaging parameters are included in supplementary materials. Image analysis for data recorded on the Cytation 5 was conducted using the BioTek Gen5 software. For time-lapse microscopy, for each sample, a region of interest (ROI) was identified and fluorescence tracked over a 24 h period.

## Results

### *S. aureus* invasion of human lung epithelial cells is dependent on circadian phase

To investigate the role of the circadian clock in the invasion of *S. aureus*, A549 human lung adenocarcinoma cells were synchronized using forskolin (FSK). A549 cells are well known to contain a functional clock^29^ and FSK is a well-known cAMP inducer commonly employed for cellular clock synchronization^19^. We confirmed circadian oscillations in the A549 cells synchronised with FSK by creating and recording from an A549;*Per2-luc* reporter cell line (Figure S2A).

Epithelial cells were synchronised by forskolin in 6-hourly intervals over a 24-h period and subsequently infected with *S. aureus* for 1 h. In addition, unsynchronised A549 cells were infected (Figure 1A). Notably, compared to unsynchronised cells, a higher number of intracellular bacteria was observed in synchronised cells after 12 h before infection, as measured by colony forming units (CFU), whereas a significantly lower uptake was detected in cells synchronised 24 h before infection. These findings suggest that the phase of the cellular circadian clock modulates the uptake of *S. aureus* (Figure 1B). Non-linear curve fitting using COSINOR analysis suggested a significant rhythmicity with and approximately 3-fold change in colony-forming units (CFU) between the 12 h and 24 h samples (sum-of-squares F-test, *P* = 0.0001, Figure 1C).

To verify this result and visualize intracellular bacterial foci, confocal microscopy was performed on fixed samples infected at different circadian phases as above. We found a higher number of intracellular bacteria within cells synchronised 12 h before infection compared to those 24 h before. Quantification of these samples showed on average of ∼3 bacteria per mammalian cell 12 h after FSK synchronisation, while this decreases at 24 h post FSK to 1.5 bacteria per cell (Figure 1E), reinforcing the notion that the host’s circadian clock may play a modulating role in bacterial uptake with an amplitude of 2-3-fold.

We further monitored the intracellular growth kinetics of *S. aureus* in the presence of 5 µg/ml lysostaphin, which kills extracellular bacteria (Figure S1A) for 12 h after infection of A549 cells using fluorescence time-lapse imaging. As above, synchronised A549 cells were infected at four distinct circadian phases. Consistent with results for invasion, our data showed a steeper increase in bacterial foci in the earlier time (6 h and 12 h) and a longer time to reach peak intracellular bacterial load (Figure S1B). Together our findings highlight a potential role for the circadian clock in modulating mammalian host cell factors for *S. aureus* cell invasion.

### Rhythmic *S. aureus* invasion is lost in clock-disrupted human epithelial cells

To investigate if the observed changes are driven by the canonical circadian clock, we generated clock-disrupted A549 cells by targeting the transcription factor BMAL1, an essential core circadian clock gene^1,30^. We generated *BMAL1* knockdown (*shBMAL1*) A549 cells using a lentiviral transduction system. Successfully transfected cells were selected based on puromycin resistance conferred by the presence of the knockdown-construct and confirmed by eGFP expression (Figure S2B). Downregulation of BMAL1 protein expression was confirmed via western blot analysis (Figure 2A). Quantitative densitometric analysis of the blots revealed a 50% reduction in BMAL1 protein levels in knockdown cells compared to parental A549 cells (Figure 2A). Gene expression profiling with RNA-Seq also confirmed downregulation of *BMAL1* mRNA and pathway analysis suggested a disruption of the KEGG pathway for “circadian rhythm” (Figure S2C).

**Figure 2.**
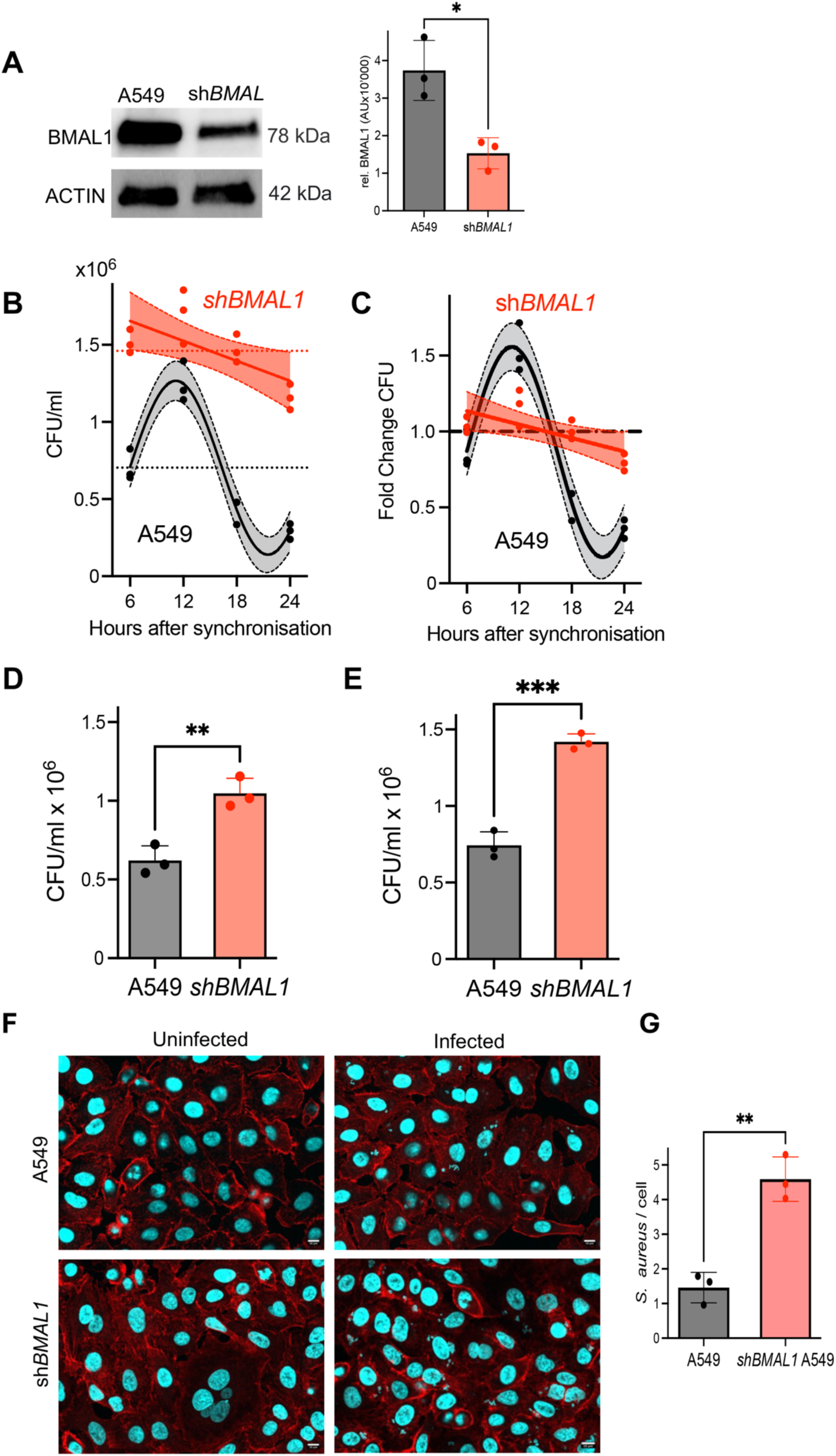
The canonical circadian clock is necessary for rhythm of S. aureus invasion, with clock-disrupted cells more permissive for S. aureus invasion. **A)** Western blot of parental and *shBMAL1* A549 cell lysates confirms BMAL1 knockdown. Quantitation of band intensities relative to ACTIN showed an approximately 2-fold reduction, N = 3, \**P* = 0.0166, unpaired t-test. Time-series of *S. aureus* invasion in parental (black: A549) and *shBMAL1* (red: *shBMAL1*) A549 cells performed as described in Figure 1A showing (**B**) bacterial colony counts (CFU/ml) and (**C**) fold change relative to the unsynchronized control. Dotted lines on graphs show the level of infection of unsynchronized control cells of each cell line. Best fit for COSINOR or linear model show depending on sum-of-squares F-Test as described in Methods section. Unsynchronised parental and *shBMAL1* A549 cells with no synchronization were infected with *S. aureus* JE2 for 1 h, extracellular bacteria removed and cells further incubated for 24 h in the presence of lysostaphin (2 μg/ml). Intracellular bacterial counts after (**D**) 1 h and (**E**) 24 h post infection. N = 3, mean+/-SD, \*\**P* < 0.01, \*\*\**P* < 0.001, unpaired t-test. (**F**) Confocal images from infected and uninfected cells 1 h after infection, Red- actin stained by Phalloidin Alexa fluor 647, Blue- bacterial DNA and cell nuclei stained by DAPI. Scale bar = 10 μm. Single Z-sections are shown (**G**) Quantification of intracellular bacteria relative to number of mammalian cells from confocal images from 3 independent experiments, N = 3, *P* = 0.0022, unpaired t-test.

We then tested the clock disrupted *shBMAL1* A549 cells in the infection assay with *S. aureus* as described above. Unlike parental cells, which again exhibited a circadian phase-dependent rhythm of bacterial invasion, a temporal invasion pattern was abolished in *shBMAL1* A549 cells (Figure 2C,D). Non-linear COSINOR fit the data statistically worse than a linear model for *shBMAL1* cells (sum-of-squares F-test, *P* = 0.0151; Figure 2C,D). This observation strongly suggests the circadian clock, for which *BMAL1* is a critical regulator, is necessary for *S. aureus* invasion rhythms in A549 cells.

Interestingly, our data in *shBMAL1* A549 cells suggests an increase in *S. aureus* invasion irrespective of time after synchronization. Quantitative CFU analysis showed that the uptake in knockdown cells was approximately 1.7-fold higher than in parental A549 cells (Figure 2E). Furthermore, intracellular bacterial survival after 24 hours was also higher for *shBMAL1* cells (Figure 2F). To visualize intracellular bacteria, we performed confocal microscopy on fixed mammalian cells. Consistent with the CFU data, microscopy images confirmed a greater number of intracellular bacteria in sh*BMAL1* cells compared to the parental line (Figure 2G,H). The significantly higher bacterial burden in *BMAL1* knockdown cells highlights its role as a regulator of host-pathogen interactions, potentially through pathways that influence cellular processes such as bacterial recognition, adhesion, or uptake.

### Effects of sh*BMAL1* knockdown on the lung epithelial cell transcriptome

To investigate the potential mechanisms underlying the increased bacterial uptake after infection with *S. aureus* 12 h versus 24 h post synchronisation and the time-invariant increase in sh*BMAL1* knockdown cells, we sequenced the transcriptome of parental and sh*BMAL1* lines at 12 h and 24 h post synchronisation. The resulting principal component analysis indicated good separation by cell line (significant correlation with PC1, *P* = 9.75e-07, Figure 3A). First, we considered the differences between the two sequenced cell lines by pooling all samples from the two experimental time points (i.e., 12 and 24 hours after synchronisation). DESeq2 analysis identified a total of 3693 differentially expressed genes (DEGs, 1773 up and 1920 downregulated) in shBMAL1 knockdown vs. parental A549 cells. Furthermore, 29 and three genes were significantly down or up-regulated, respectively, at 12 h compared to 24 h post synchronisation, and no DEGs were found for the interaction term of cell line and timing. Given the low number of samples, this was not unsurprising. However, pathway analysis using PGSEA indicated that 17 KEGG pathways were significantly enriched (Figure 3B). Reassuringly, “circadian rhythms” was significantly less active in the knockdown cells (Figure 3B, Figure S2D). Of the other 16 pathways, at least half have some relevance with infection and inflammation processes. Interestingly, “*S. aureus* infection” was among the pathways significantly enriched (Figure 3B).

**Figure 3.**
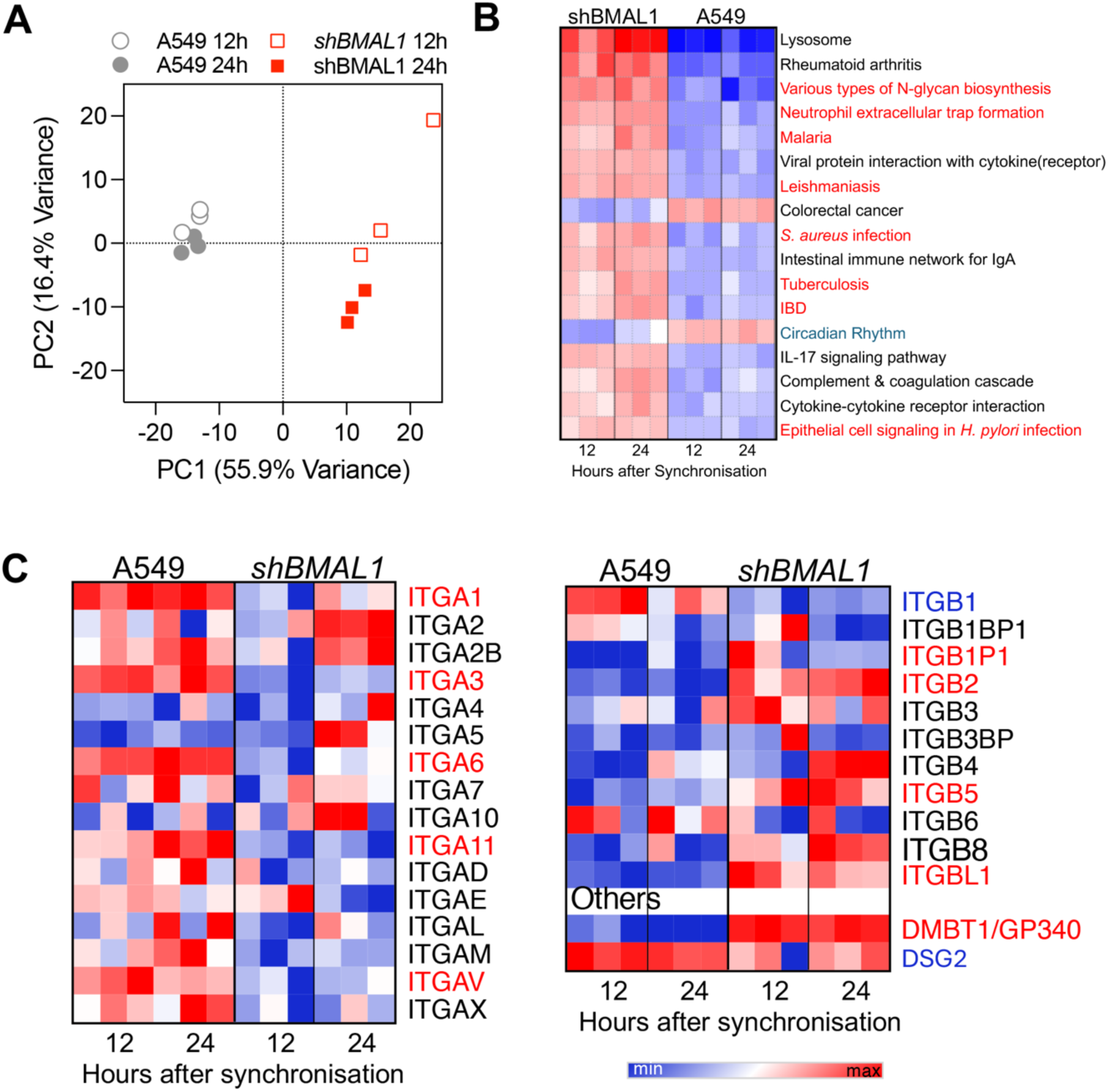
*BMAL1* knockdown cells exhibit significant changes in gene expression. (**A**) PCA plot comparing the variance in gene expression by RNA-Seq between parental and shBMAL1 A549 cells collected at 12 h and 24 h after synchronization. (**B**) Significantly regulated KEGG pathways by PGPSEA analysis. Pathways in red font are infection and blue font circadian rhythm related. (**C**) Heatmap showing gene expression profiles for genes encoding integrin A (ITGA) and B (ITGB) subunits and other receptors (DMBT1/GP340) involved in *S. aureus* uptake profiles in parental and *shBMAL1* A549 cells. Names of differentially expressed genes in red font.

Contrary to expectations, we did not observe consistent changes in the expression of genes encoding integrins α5 and β1, the receptors to which *S. aureus* adhesins fibronectin binding proteins, Fnbp A and Fnbp B, bind to for the invasion of epithelial cells^31,32^. The gene encoding for *ITGA5* was not significantly changed, and *ITGB1* was even significantly downregulated (Figure 3C). However, when we systematically assessed all integrin isoforms and other extracellular proteins known to be involved in *S. aureus* invasion, we found several other integrin isoforms, e.g., *ITGB2* and *ITGB5* upregulated in the *shBMAL1* A549 cells (Figure 3C).

Another differentially expressed gene which was previously recognised to be important for *S. aureus* uptake is GP340^33,34^ (Figure 3C). When we compared GP340 coding gene (*DMBT1*) expression at 12 vs. 24 hours after synchronisation, there was only a subtle difference (Figure 4A). In line with our above observations, however, we observed a substantial upregulation of the *GP340* coding gene in *BMAL1* knockdown cells (Figure 4A). We further characterised GP340 protein dynamics using flow cytometry on non-permeabilised live cells and confirmed a 4-fold higher GP340 expression in *shBMAL1* compared to parental A549 cells (Figure 4B). Reassuringly a higher abundance of GP340 positive cells at 12 hours after synchronisation compared to 18 and 24 hours after synchronisation (Figure 4 C,D, Kruskal-Wallis, *P* = 0.0029, adj. *P* of Dunn’s post-hoc 12 vs. 18 *P* = 0.0033, 12 vs 24 *P* = 0.0075).The COSINOR analysis suggested a rhythmic pattern with 24-hour period was the better fit for the data (COSINOR vs. line: *P* = 0.0084). The bacterial ligand to GP340 is the *S. aureus SraP* protein, which has been reported to facilitate bacterial uptake^33,34^. Thus, *GP340* may be important in the circadian modulation of *S. aureus* uptake.

**Figure 4.**
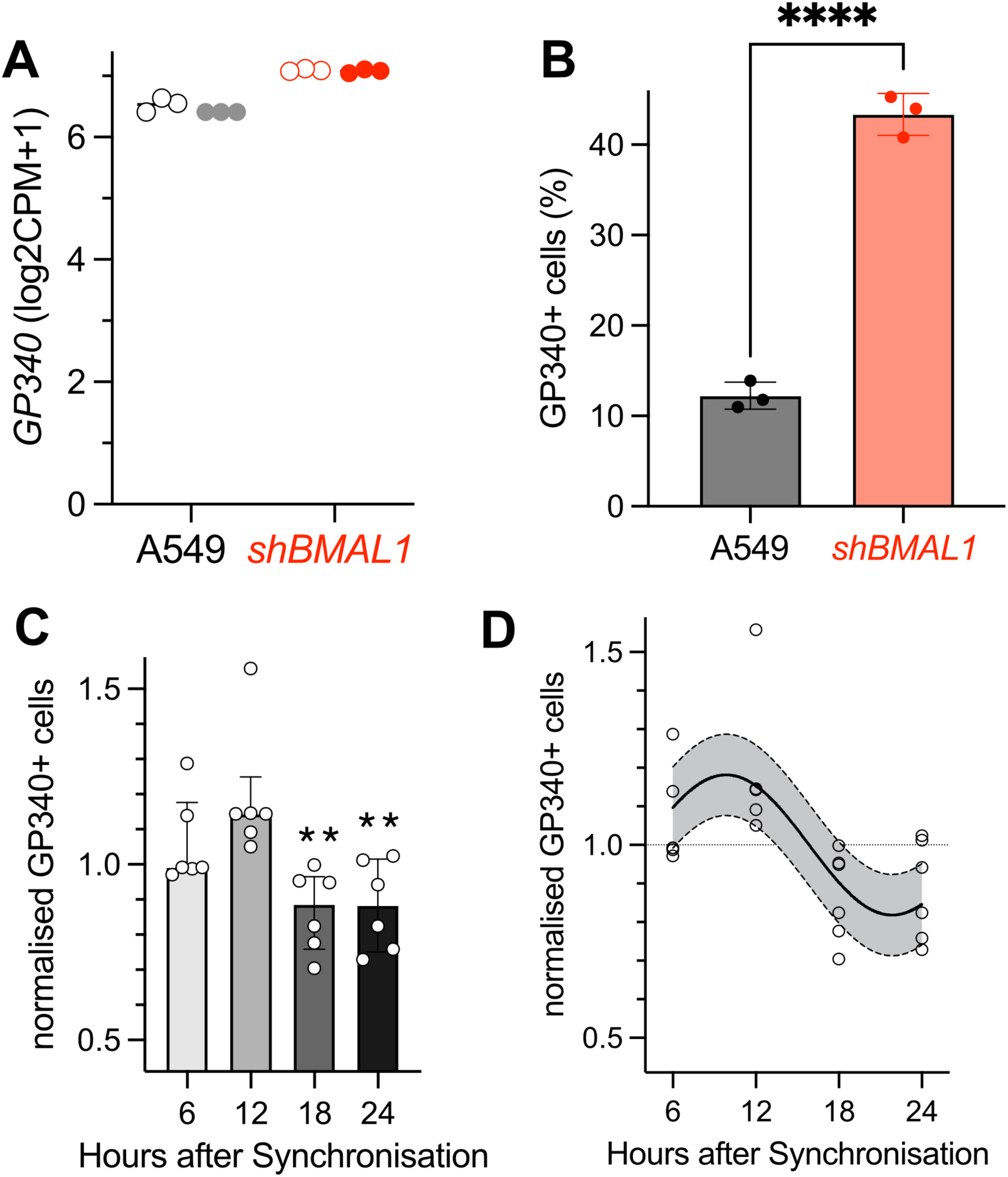
GP340 protein expression exhibits a daily rhythm in A549 cells and is increase in *BMAL1* knockdown cells. (**A**) Normalised gene expression of GP340 at 12 h (open circles) and 24 h (closed circles) in parental A549 and *shBMAL1* knockdown cells. (**B**) Proportion of GP340 positive cells assessed by flow cytometry is increased (t-test, *P* <0.0001) in *shBMAL1* compared to A549 parental line. (**C&D**) In A549 cells, GP340 protein expression levels were higher at 12 hours after FSK synchronization compared to 18 and 24 hours (Kruskal-Wallis test, *P* = 0.0029. Dunn’s test: 12 vs. 18: *P* = 0.0033 and 12 vs. 24: *P* =0.0075). Expression levels were normalized to median number of GP340+ cells. Model comparison suggested COSINOR to be the preferred fit when compared to linear (*P* = 0.0084, Best fit for COSINOR or linear model show depending on sum-of-squares F-Test as described in Methods section).

### The *S. aureus* adhesin *SraP* mediates higher bacterial uptake by *shBMAL1* cells

To directly test the role of *GP340* in rhythmic *S. aureus* invasion, we compared the internalisation of an *S. aureus sraP* mutant with the WT JE2 strain used above in both, the parental and *shBMAL1* A549 cells. Consistent with the data shown in Figures 2 and 3, the uptake of *S. aureus* JE2 by sh*BMAL1* knockdown cells was significantly increased compared to parental A549 cells, however, uptake of the *sraP* mutants in *shBMAL1* was similar to that by the parental A549 cells (Figure 5A). Importantly, in both cell lines, and consistent with Figures 1 and 2, JE2 but not s*raP* mutants showed rhythmic invasion in A549 cells. As expected, both strains did not show rhythmic invasion into *shBMAL1* A549 cells (Figure 5B), but a much higher invasion in non-synchronised cells (Figure 5A).

**Figure 5.**
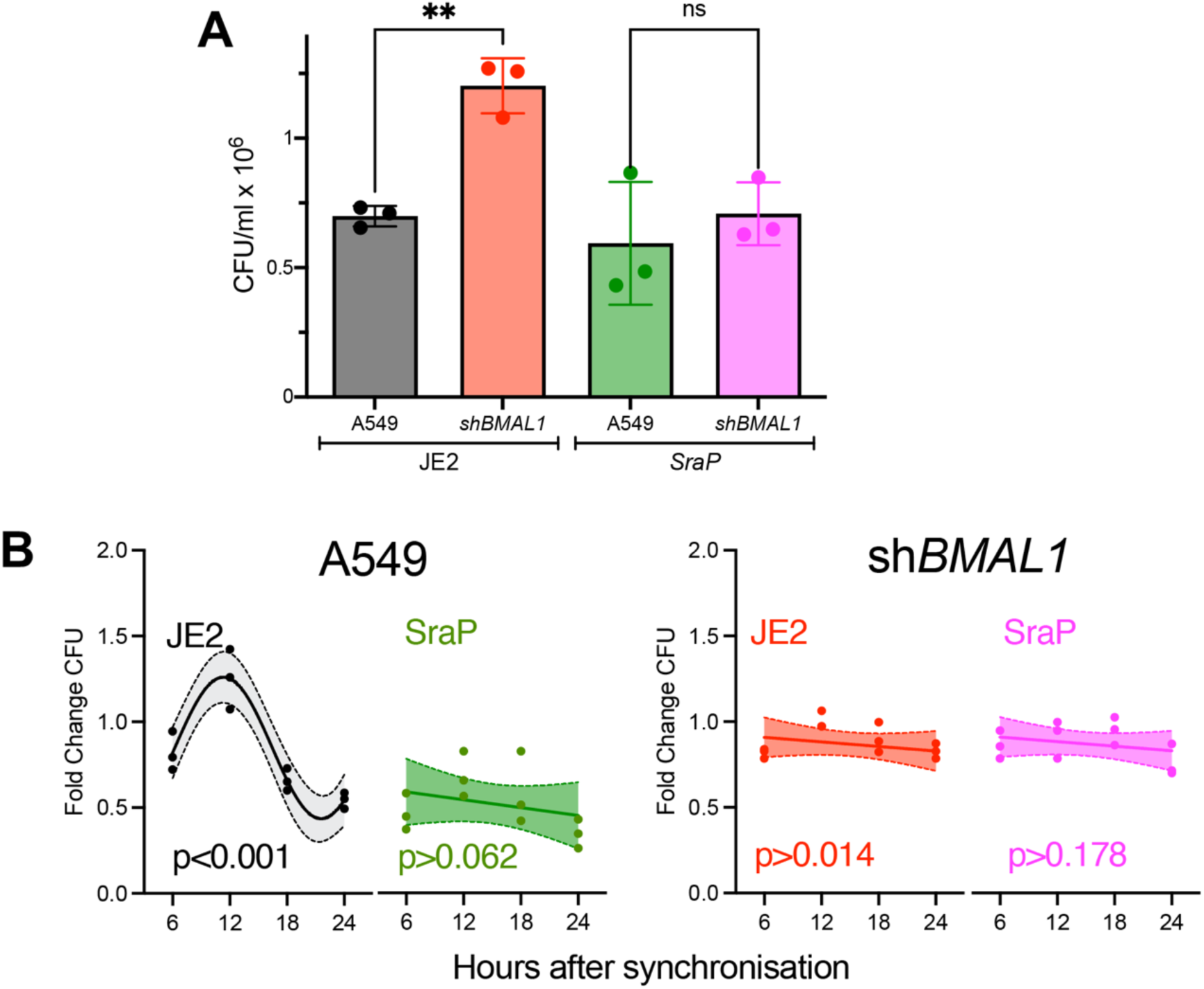
GP340 ligand SraP mediates the rhythmic uptake of *S. aureus* in lung epithelial cells. (**A**) Non-synchronized parental and *shBMAL1* A549 cells were infected with WT *S. aureus* (JE2) or *S. aureus sraP* transposon mutant for 1 h, and intracellular bacterial numbers as measured by CFU were compared, ** *P* < 0.01 by an unpaired t-test (**B**) Infection of *S. aureus* JE2 and *sraP* mutant at four times after synchronization of cells with FSK as in Figure 1. The fold changes of CFU for each strain and time compared to the respective control of unsynchronized cells of hat strain are plotted. COSINOR and linear model fit are compared by extra sum of squares F-test and better fit is plotted as described in Methods section. N = 3.

Additionally, we infected both cell types with transposon mutants lacking FnbpA and FnbpB), key *S. aureus* adhesins/invasins. We observed that while the uptake of these mutants was also higher in *shBMAL1* when compared to parental A549 cells, there was no difference when we compared fold changes relative to WT JE2 (Figure S3B). Thus, in contrast to other adhesins tested, our data suggest that *S. aureus* SraP mediate rhythmic bacterial uptake through GP340 and the observed increase in *BMAL1* knockdown cells.

## Discussion

The ability of *S. aureus* to invade and survive within mammalian cells has been attributed to bacterial persistence, antibiotic resistance and recurrent infections. Although *S. aureus* factors that enable cell invasion have been well studied, the host cellular pathways that mediate this are less understood. Circadian rhythms have been demonstrated to influence several host immune responses during infection with some bacteria exhibiting time-of-day differences in infecting flies and mice^10^. However, at the cellular level, the role of the circadian clock in controlling pathogen invasion or intracellular survival is not known. Here we provide evidence for the role of the mammalian circadian clock in modulating invasion of *Staphylococcus aureus* in human lung epithelial A549 cells. We found significant temporal variations in bacterial uptake, a peak in invasion 12 h after forskolin (FSK) synchronisation of the A549 cells and a marked decrease 24 h post synchronisation. Disrupting the clock through *shBMAL1* knockdown not only abrogated the rhythm in invasion, but also increased the bacterial load, highlighting the influence of the canonical circadian clock in host-pathogen interactions at a cellular level. Notably, using transcriptomic and protein expression analysis, we identified a significant rhythm in GP340 protein abundance and upregulation of *GP340* in the clock-disrupted *shBMAL1* A549 cells. GP340 is known to be a key factor for the uptake of *S. aureus* by epithelial cells through the *S. aureus* adhesin SraP^33–35^, and our data suggest that GP340 may mediate the rhythmic uptake of bacteria.

Circadian regulation of immune responses is well documented, with evidence suggesting that various immune processes, including cytokine production and phagocytosis, exhibit circadian variations^36^. Colonisation by bacterial pathogens including *Salmonella typhimurium, Streptococcus pneumoniae and S. aureus* have been shown to be impacted by the time of the day, primarily driven by clock-dependent immune responses to infection^37^. As intracellular survival of many of these pathogens have been reported to be an important feature of bacterial pathogenesis, our findings could be relevant for multiple intracellular pathogens. Our findings extend this knowledge by demonstrating that synchronization with FSK not only impacts the invasion of *S. aureus* but also alters the intracellular responses to bacterial challenge.

*BMAL1* has a key role in the regulation of circadian rhythms^30^. This complex also activates the transcription of negative regulators, which feedback to inhibit BMAL1-CLOCK activity, ensuring tightly regulated 24-hour rhythms in gene expression^38^. It has been shown that the circadian transcription factor BMAL1 modulates hepatitis C virus (HCV) entry into hepatocytes by regulating the expression of the viral receptors *Cd81* and *Claudin-1*^39^. Pharmacological inhibition of *Bmal1* by REV-ERB agonists reduced *Cd81* and *Claudin-1* expression and inhibited viral entry in cells^39^. Similarly, this was found to down-regulated *angiotensin-converting enzyme 2* expression, which is essential for the entry and replication of SARS-CoV-2 in lung epithelial cells^40^. This highlights the importance of circadian processes for the treatment of viral infection. In addition, various parasitic infections are also linked with regulation of circadian clock^12^. Importantly, it has been reported that the complete loss of circadian regulation in *Bmal1* knockout macrophages alters the structure of actin cytoskeleton, which is coupled with a significant increase in *S. pneumoniae* burden compared to WT cells, emphasizing a role for *Bmal1* in limiting bacterial uptake and proliferation^41^.

Our data demonstrated increased bacterial uptake in *BMAL1* knockdown cells, which occurred independent of the uptake effects of FSK-mediated circadian synchronization of the population. Thus, the temporal modulation in bacterial uptake cannot be explained by the effect of FSK on cAMP induction per se but establishes a role for *BMAL1*-regulated processes in host susceptibility to *S. aureus* invasion. These observations align with the report that BMAL1 deficiency impacts immune cell function and can modulate inflammatory responses^42^.

Microbial surface components recognizing adhesive matrix molecules (MSCRAMMs) are most prominent surface proteins expressed by staphylococci, which facilitate adhesion to the extracellular matrix (ECM) components such as collagen, fibrinogen, and fibronectin (Fn)^43^. Although, the fibronectin-binding protein (FnBP)–fibronectin (Fn)–α5β1 integrin pathway is the primary route for *S. aureus* internalization^32,44^, we did not find upregulation of the main integrins α5β1 in parental vs. *shBMAL1* A549 cells. Although *fnbA* and *fnbB* mutants showed reduced uptake in parental cells, they showed higher invasion in *shBMAL1* A549 cells, suggesting involvement of secondary mechanisms in bacterial uptake. We have not used a double mutant lacking both *fnbA* and *fnbB* in this study, so a role of Fnb proteins cannot be fully ruled out. As we observed differential regulation of several integrin isoforms in the RNA-seq results, it is possible that these other isoforms are potentially involved in bacterial uptake. The specific roles of such isoforms in *S. aureus* uptake, however, have not been extensively studied and their potential involvement in bacterial uptake warrants further investigation. As BMAL1 can impact multiple cell signalling pathways, these may also contribute to the increased bacterial uptake^45^.

Fibronectin-independent mechanisms are known to mediate *S. aureus* entry and involve distinct surface proteins such as serine-aspartate repeat-containing protein D (SdrD), clumping factor A (ClfA), autolysin (Atl), and serine-rich adhesin for platelets (SraP) ^43^. SdrD directly binds Desmoglein 1 (encoded by *DSG1*)^46^, ClfA interacts with host cells either directly or via fibrinogen bridges^47^ and Atl associates with heat shock cognate protein 70 (*HSPA8*)^48^, while SraP binds to the salivary scavenger receptor GP340^33,34^. Interestingly, we found a notable rhythm in GP340 protein expression. Circadian disrupted shB*MAL1* knockdown cells, showed increase GP340 gene and protein expression and a reduced invasion of *sraP* mutant strains, suggesting a role for the SraP-GP340 pathway in the circadian modulation of bacterial invasion. SraP, a member of a family of serine rich glycoproteins encoded by Gram positive bacteria including *Streptococcus spp*^35^, was shown to mediate adhesion to epithelial cells through its L-lectin binding module^33^. It was also recently reported to modulate macrophage apoptosis and inflammation, indicating its role in *S. aureus*-immune cell interactions^49^. While GP340 is implicated as a receptor in clock-dependent *S. aureus* invasion, it could serve as a co-receptor with other integrins, mediating alternative mechanisms of invasion.

Despite the robust findings of this study, our data are from an epithelial cell line (A549) which may limit the generalizability of the results across different cell types and physiological contexts. Future studies should examine the impact of circadian rhythms on *S. aureus* invasion in a broader range of epithelial and immune cell types, in particular immune cells to link these findings better to previous studies on circadian effects in immune responses. Additionally, while our study provides insights into the role of *BMAL1* in regulating bacterial invasion, the specific downstream signalling pathways and molecular mechanisms to understand how circadian rhythms influence the expression of key receptors involved in bacterial invasion remain to be elucidated.

The implications of the findings from this study extend to the potential for developing therapeutic strategies that target circadian rhythms to improve the timing and efficacy of antimicrobial treatments. For instance, studies have shown that administering antibiotics in alignment with the host circadian cycle can enhance their effectiveness and reduce side effects^50^. Recent studies have reported intestinal clocks to control the gut microbiota composition and function^51^, with enteric bacteria also responding to gastrointestinal hormones like melatonin^52^. While circadian clocks have been only recently described in bacteria^53,54^, it would be interesting to study if bacterial pathogens have circadian rhythms and if these pathogen clocks interact with mammalian clocks. Such interacting clocks have been described elsewhere, for example in legumes and rhizobia^55^.

In conclusion, our study highlights the critical role of circadian rhythms in modulating the dynamics of *S. aureus* invasion in epithelial cells. Overall, our results highlight the critical interplay between circadian biology and microbial pathogenesis, suggesting that manipulation of circadian pathways may provide new avenues for innovative strategies aimed at preventing and treating bacterial infections.

## Acknowledgements

We are grateful for funding from the Warwick Research Development Fund (RD and MU) and a Monash Warwick Alliance fellowship to PA (MU). Anglo American Platinum is gratefully acknowledged for funding work in RDs group, and James Jarrold for help with sample collection. Zijin Zhang is thanked for generating the A549 Per2-luc bioluminescence data. We acknowledge use of the flow cytometry equipment and the support provided by the Bio-Analytical Shared Resource Laboratories within the University of Warwick. We thank Vadim Vasilyev for advice on the RNASeq analysis.

## Author contributions

P.A. designed and carried out the experimental work, analysed the resulting data and co-wrote the manuscript.

G.C., S.N. and J.G. contributed to the experimental design, carried out experimental work and contributed to data analysis and writing of the manuscript.

R.D. and M.U. jointly supervised the work, and contributed to formal analysis, writing of the manuscript and acquisition of funding.

## Data availability

Summary data are depicted in the figures. Raw data are available from the corresponding authors upon reasonable request. RNA-Seq data are available at the ELIXIR database ArrayExpress under accession number E-MTAB-15036.

