## Supplementary Figures for "The cellular mammalian clock regulates *Staphylococcus aureus* invasion in epithelial cells"

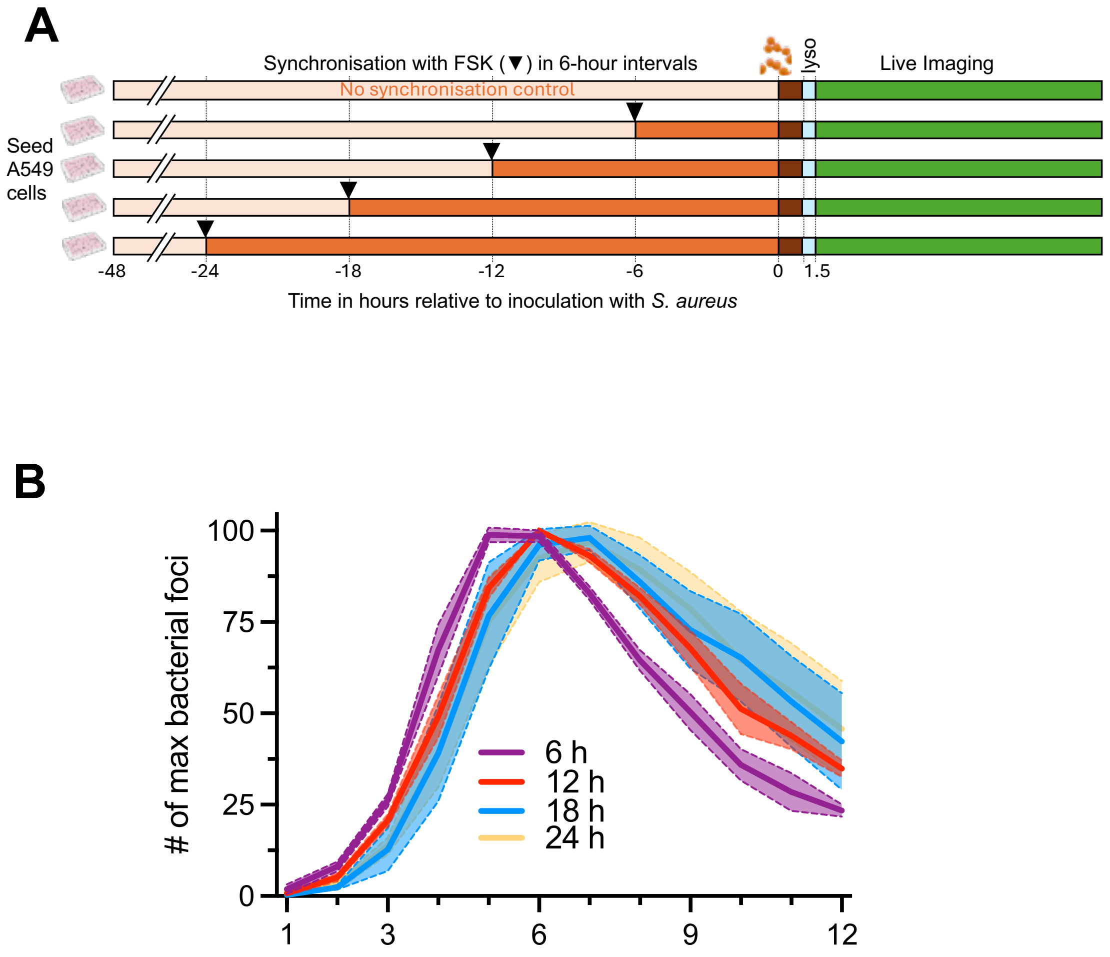


**Figure S1**. **Bacterial growth kinetics is delayed at 24 h compared to 12 h after synchronisation. (A)** A549 epithelial cells were synchronized with FSK every 6 hours over 24 hours period and infected with *S. aureus* for 1 h, extracellular bacteria were removed by gentamycin/lysostaphin treatment and cells imaged in presence of lysostaphin for 12 h. (B) Number of intracellular *S. aureus-GFP* foci plotted as hourly running average.

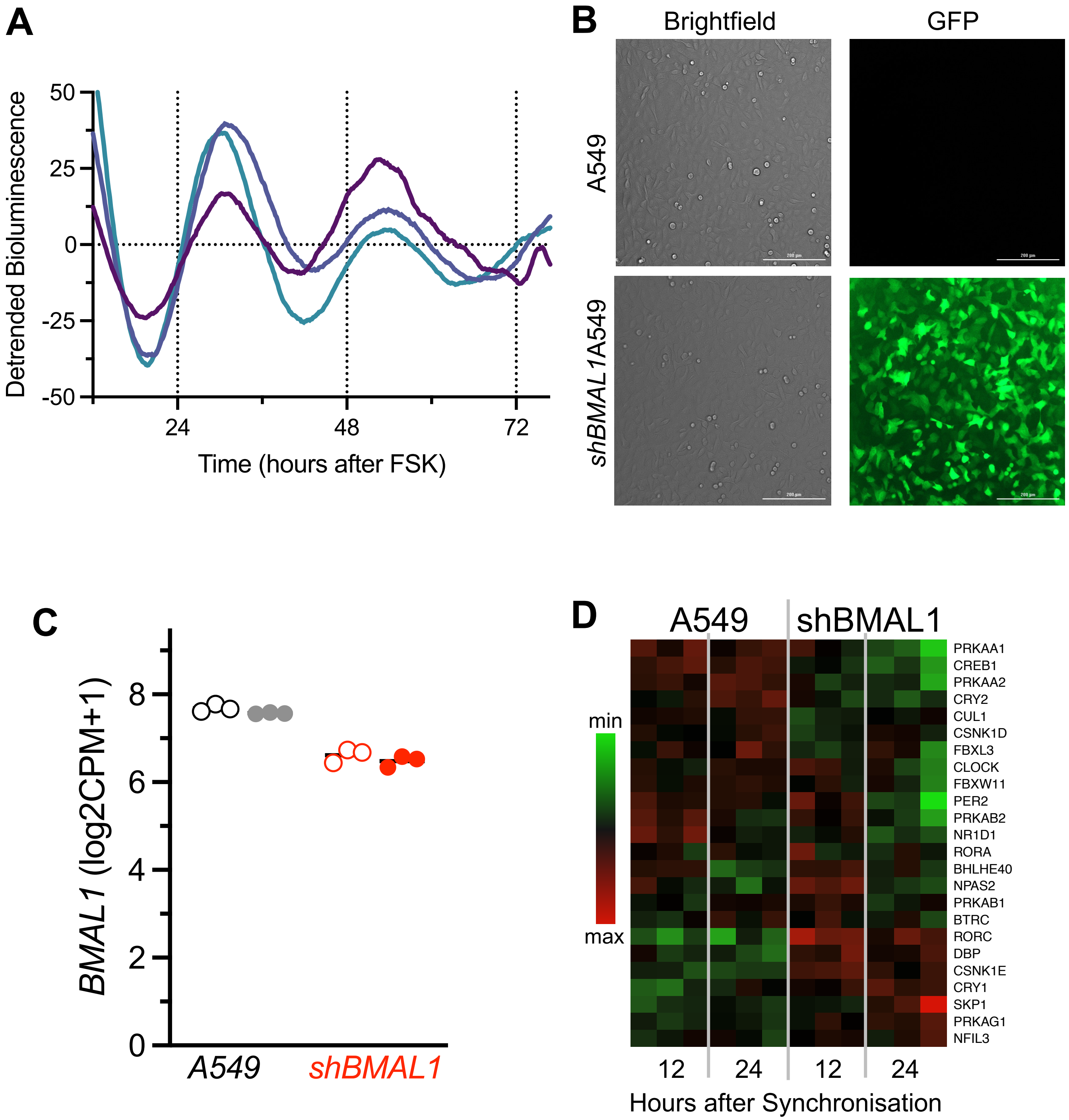


**Figure S2**. **A549 circadian properties and knockdown of BMAL1 gene expression**. (A) Three replicate dishes of FSK synchronised A549;Per2-luc cells show consistent circadian oscillations in bioluminescence. For this, A549 reporter cells were generated as described for shBMAL1 cells in the methods using the Per2-luc viral construct and bioluminescence recording methods as described previously (Zhang et al. 2017 doi: 10.1080/15384101.2017.1387695). Each line represents one replicate. (B) Generation of stable *shBMAL1* cell line expressing puromycin and eGFP. Bright field and fluorescent microscopic images of A549 and A549 shBMAL1 are shown scale bar = 200 µm **(C)** Quantification of knockdown efficiency from RNA-Seq experiment at 12h (open circles) and 24h (filled circles). Data shown as edgeR transformed (log2CPM+1) values. (**D**) Heatmap for all genes in Circadian Rhythm KEGG pathway (hsa4710) from PGSEA was significantly deregulated (N = 3, condition and cell line).

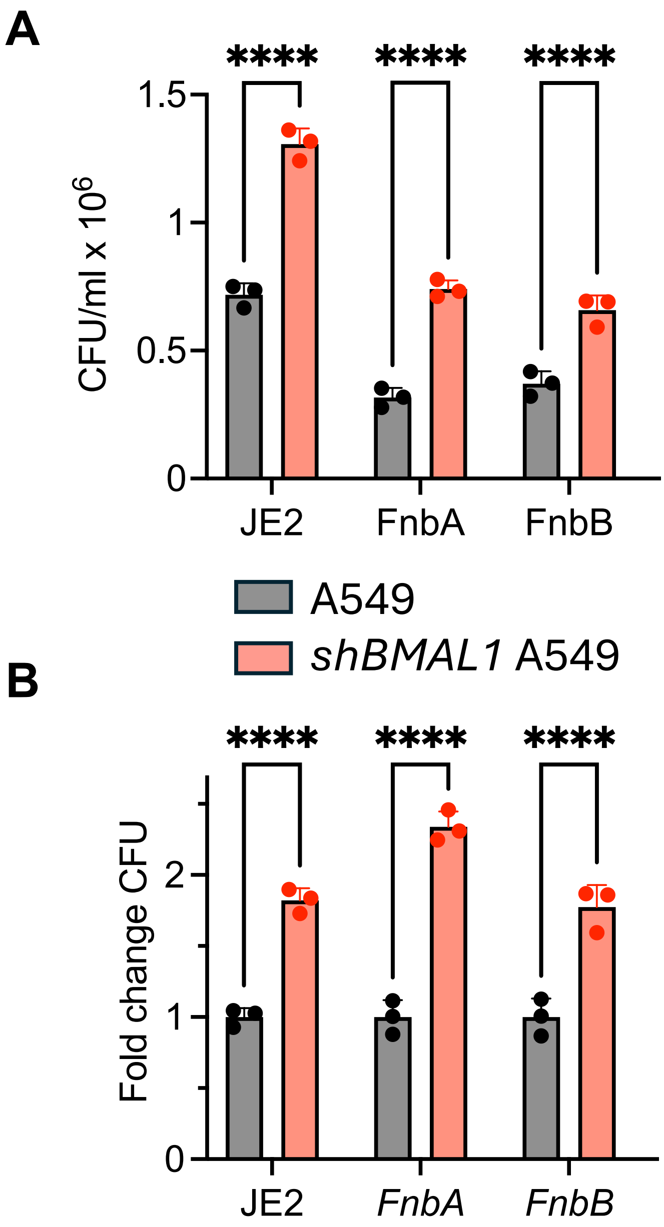


**Figure S3. Comparison of the invasion of *S. aureus* JE2 and transposon mutants of *FnbA* and *FnbB* genes.** A549 and *shBMAL1* A549 cells were infected with JE2 *S. aureus WT* or mutant strains *f*or 1 hour and intracellular bacteria were quantified by colony counts. (A) Bacterial counts (CFU) after 1 hour infection (B) Fold change in CFU calculated relative to JE2 strain. If statistically significant 2-way ANOVA result, Sidac post-hoc test given as ****$P$ < 0.0001 for strain comparison.

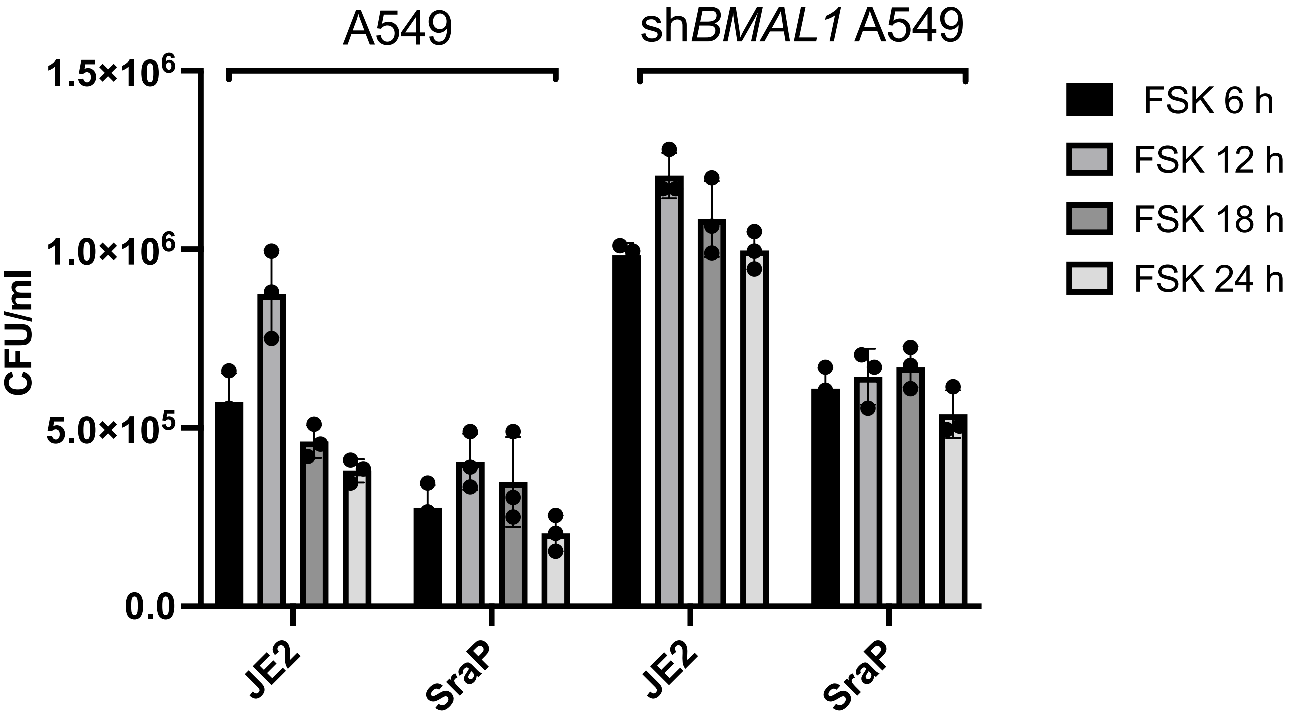


**Figure S4.** Invasion of *S. aureus* JE2 and *sraP* mutants at four times after synchronisation. A549 or shBMAL1 A549 cells were synchronized with forskolin (FSK) 6, 12, 18 and 24 h before infection with bacteria at an MOI of 10:1. CFU counts after different times of synchronization are shown. N = 3

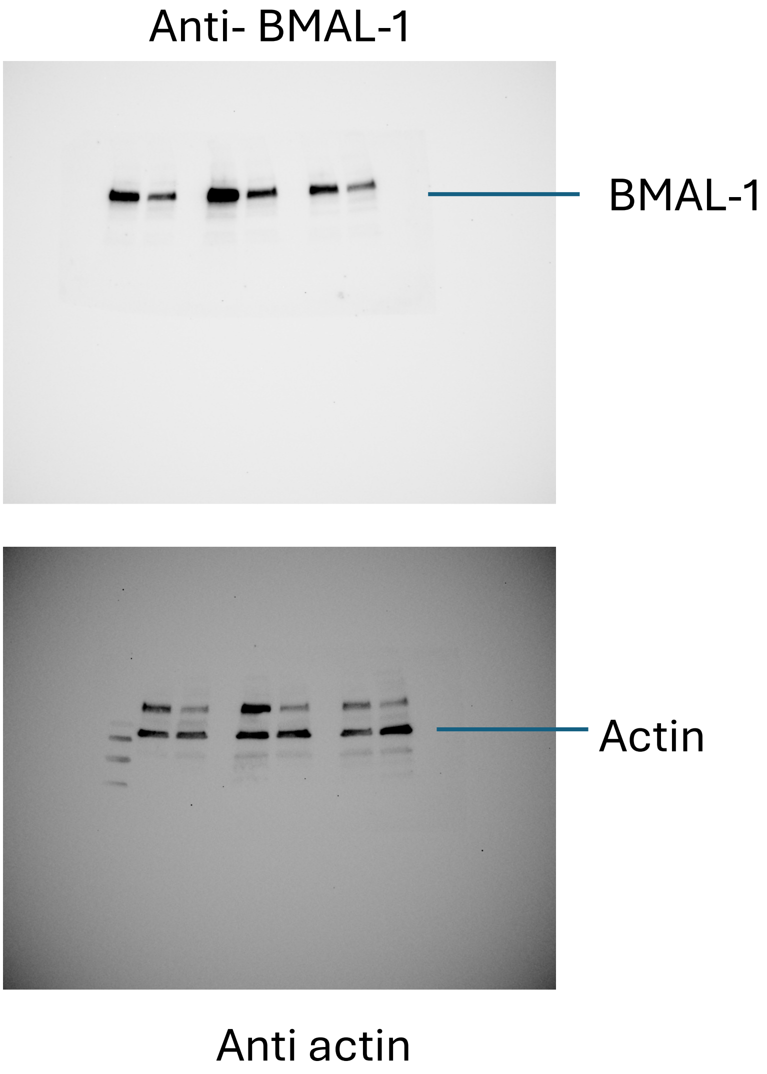


**Figure S5.** Uncropped blots shown in Figure 2A.

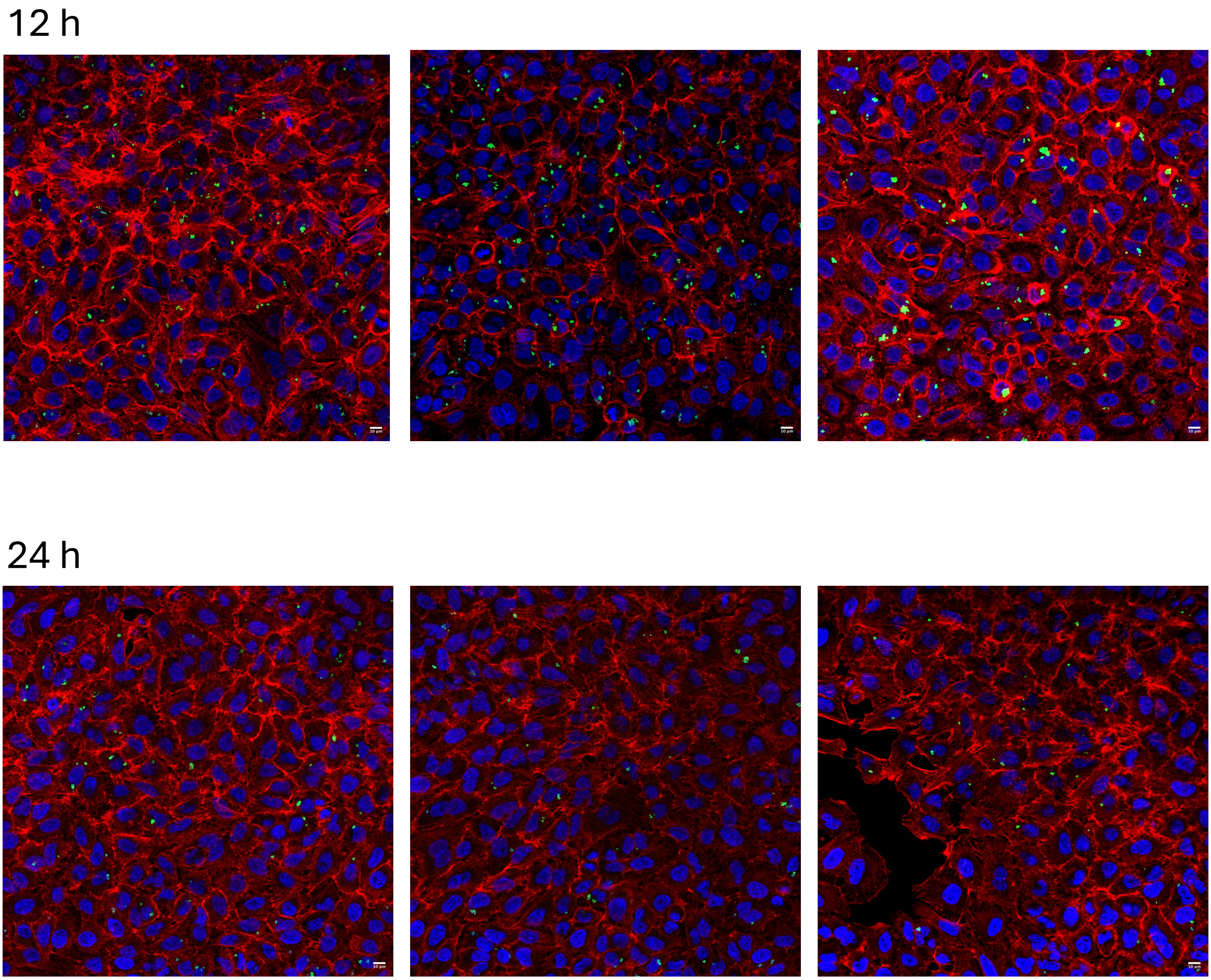


**Figure S6.** Images used in Figure 1D.

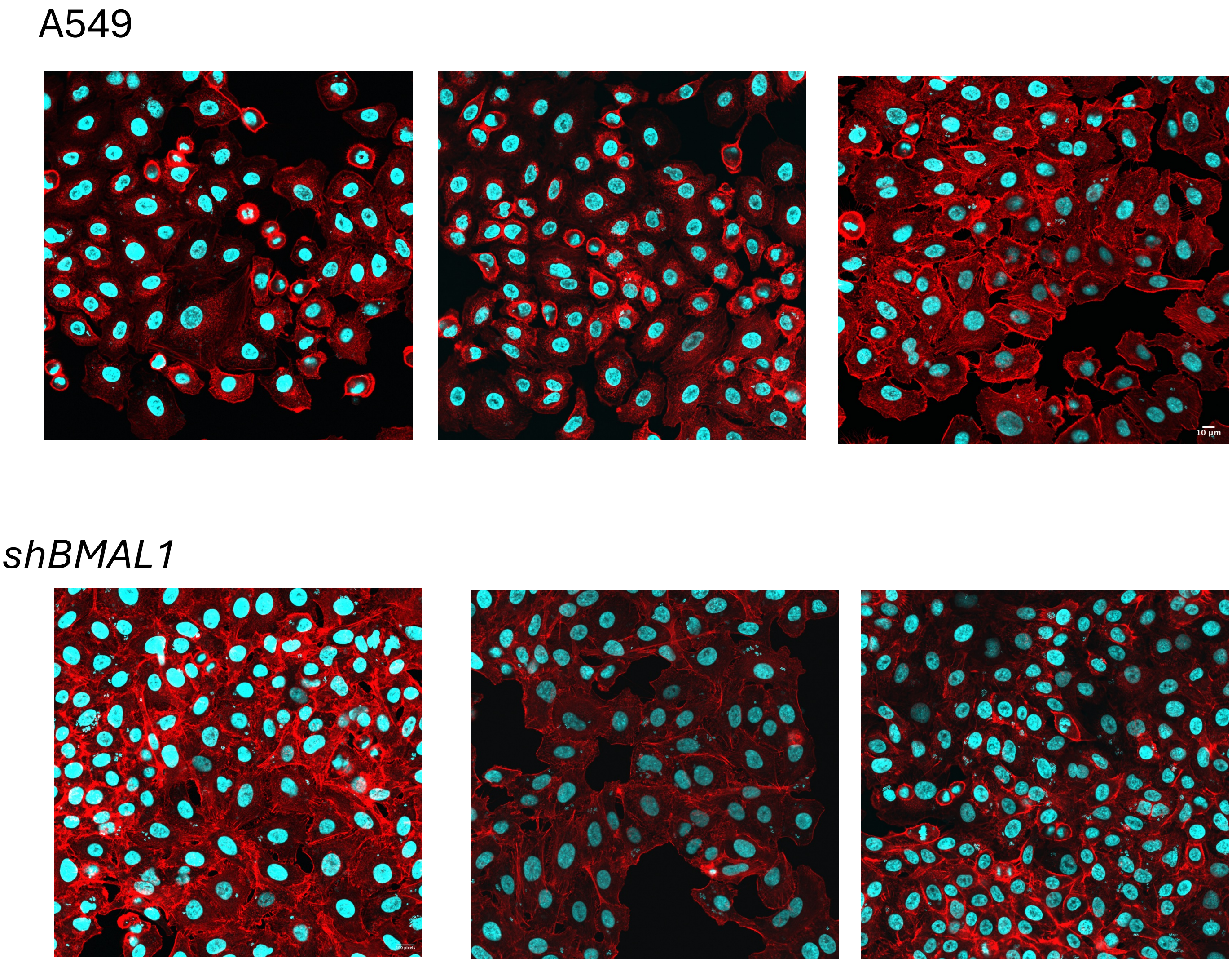


**Figure S7.** Images used for quantitation in Figure 3D.
